# Primary case inference in viral outbreaks through analysis of intra-host variant population

**DOI:** 10.1101/2020.09.18.303131

**Authors:** J. Walker Gussler, David S Campo, Zoya Dimitrova, Pavel Skums, Yury Khudyakov

## Abstract

Investigation of outbreaks to identify the primary case is crucial for the interruption and prevention of transmission of infectious diseases. These individuals may have a higher risk of participating in near future transmission events when compared to the other patients in the outbreak, so directing more transmission prevention resources towards these individuals is a priority. Genetic characterization of intra-host viral populations, although highly efficient in the identification of transmission clusters, is not as efficient in routing transmissions during outbreaks, owing to complexity of viral evolution. Here, we present a new computational framework, PYCIVO: <u>p</u>rimar<u>y</u> <u>c</u>ase <u>i</u>nference in <u>v</u>iral <u>o</u>utbreaks. This framework expands upon our earlier work in development of QUENTIN, which builds a probabilistic disease transmission tree based on simulation of evolution of intra-host hepatitis C virus (HCV) variants between cases involved in direct transmission during an outbreak. PYCIVO improves upon QUENTIN by also implementing a custom heterogeneity index which empowers PYCIVO to make the important ‘No primary case’ prediction. One or more samples, possibly including the primary case, may have not been sampled, and this designation is meant to account for these scenarios. These approaches were validated using a set of 105 sequence samples from 11 distinct HCV transmission clusters identified during outbreak investigations, in which the primary case was epidemiologically verified. Both models can detect the correct primary case in 9 out of 11 transmission clusters (81.8%). However, while QUENTIN issues erroneous predictions on the remaining 2 transmission clusters, PYCIVO issues a null output for these clusters, giving it an effective prediction accuracy of 100%. To further evaluate accuracy of the inference, we created 10 modified transmission clusters in which the primary case had been removed. In this scenario, PYCIVO was able to correctly identify that there was no primary case in 8/10 (80%) of these modified clusters. This model was validated with HCV; however, this approach may be applicable to other microbial pathogens.

A version of this software is publicly available at the following url: https://www.github.com/walkergussler/PYCIVO

## Introduction

Hepatitis C virus (HCV) infection affects nearly 3% of the world’s population and is a major cause of liver disease worldwide [1]. In the United States, HCV infection is an important public health problem, being the most common chronic blood-borne infection as well as the leading cause for liver transplantation [2]. Since 2007, HCV surpasses HIV as a cause of death in the US [3]. Outbreaks of HCV infections are associated with unsafe injection practices, drug diversion, and other exposures to blood and blood products.

RNA viruses such as HCV exist as a heterogeneous population of closely related but genetically distinct variants, known as quasispecies [4]. When a transmission event occurs between an infected person and a susceptible person, the target patient will not receive the entire intra-host HCV population from the infected individual, but only some of the variants. These variants may or may not be representative of the infected individual’s HCV population as a whole [5].

Owing to this, methodologies which rely on one consensus sequence per patient are not effective in detection of the primary case. A better alternative is to obtain a large sample of viral variants that adequately represent intra-host viral sub-populations and thus improve accuracy of genetic detection of transmissions. This is achieved through sampling intra-host variants using amplicon-based deep sequencing technology of a highly variable region of the viral genome [6].

We previously developed the Global Hepatitis Outbreak and Surveillance Technology (GHOST), a cloud-based bioinformatics suite that can infer HCV transmission clusters by analyzing intra-host HCV populations [7]. However, given a transmission cluster, GHOST lacked the functionality to identify the likely primary case: the individual who was infected the longest and has the greatest chance to have infected other persons in the outbreak. Given that the identification of the primary case can help in the interruption and prevention of outbreaks, we present PYCIVO, a model that evaluates the strength of the molecular evidence available to identify the potential primary case for any given recent transmission cluster.

## Methods

Given a transmission cluster of several samples with available sequence data, PYCIVO identifies a potential primary outbreak case using a consensus method. Two primary case *indication values* are computed for each patient in the outbreak. We call the first indication value *V_1_*; this is estimated using data from a population-based evolutionary simulator. The second indication value, named *V_2_*, is a composite heterogeneity score which is generated using an ensemble of different metrics. Identification of the likely primary outbreak case in a cluster improves with application of both values over only one of them.

### Evolutionary Simulations

To generate *V_1_*, an evolutionary distance is determined between every pair of samples in the transmission cluster and stored in a matrix *M_T_*. In order to obtain these distances, we utilize an evolutionary simulation method proposed in our earlier paper [8]. Briefly, evolutionary-based random processes are simulated over a graph *G= (V, E)*, in which the nodes are haplotypes (unique or distinct sequences), and edges exist between two haplotypes at hamming or edit distance equal to 1. In each simulation, one population is designated as a potential infector and the other as a possible infectee. The distance measure between populations is defined as the analogue of the cover time for the random process.

For this evolutionary simulator to function, it is important to reconstruct unsampled variants that were likely present between the time of transmission and the time of sampling and were linking both samples. This reconstruction is done using maximum parsimony, assuming the minimal number of mutation steps to reach from primary case to target sequences. To achieve this, we first construct a *k-step network* [6, 9-12], i.e. the union of all minimum spanning trees of the complete graph on the set of haplotypes with edge weights corresponding to hamming distance between two given sequences. Next, each edge of length r is subdivided by *r-1* vertices. These vertices represent unsampled haplotypes and form “bridges” between connected components of observed haplotypes. This procedure is outlined in figure 1. The result of the procedure is a connected graph whose edges represent single mutations and whose nodes are haplotypes of three types: those present in the potential infector, those present in the possible infectee, and simulated intermediate sequences which likely were present at some point between the time of transmission and the time of sampling. Evolutionary simulations are conducted on this constructed graph.

**Figure 1:**
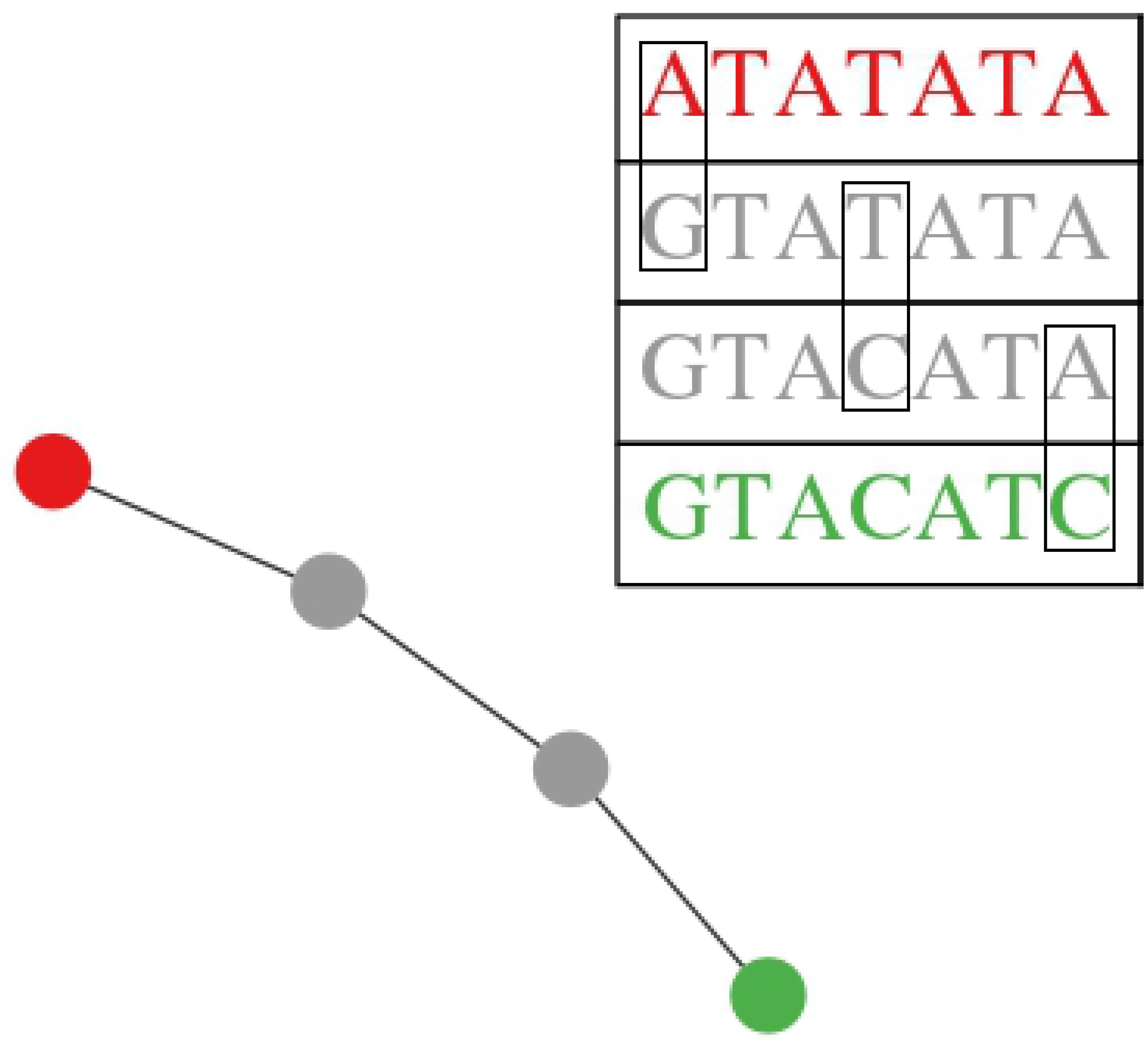
Example of a bridge reconstruction using maximum parsimony. If the red variant A were to mutate to the green variant B the gray intermediate variants would represent the most parsimonious path between these two variants.

Simulations are performed using a model from [8], which is essentially a quasispecies model with logistic growth, depicted in Figure 2.

**Figure 2:**
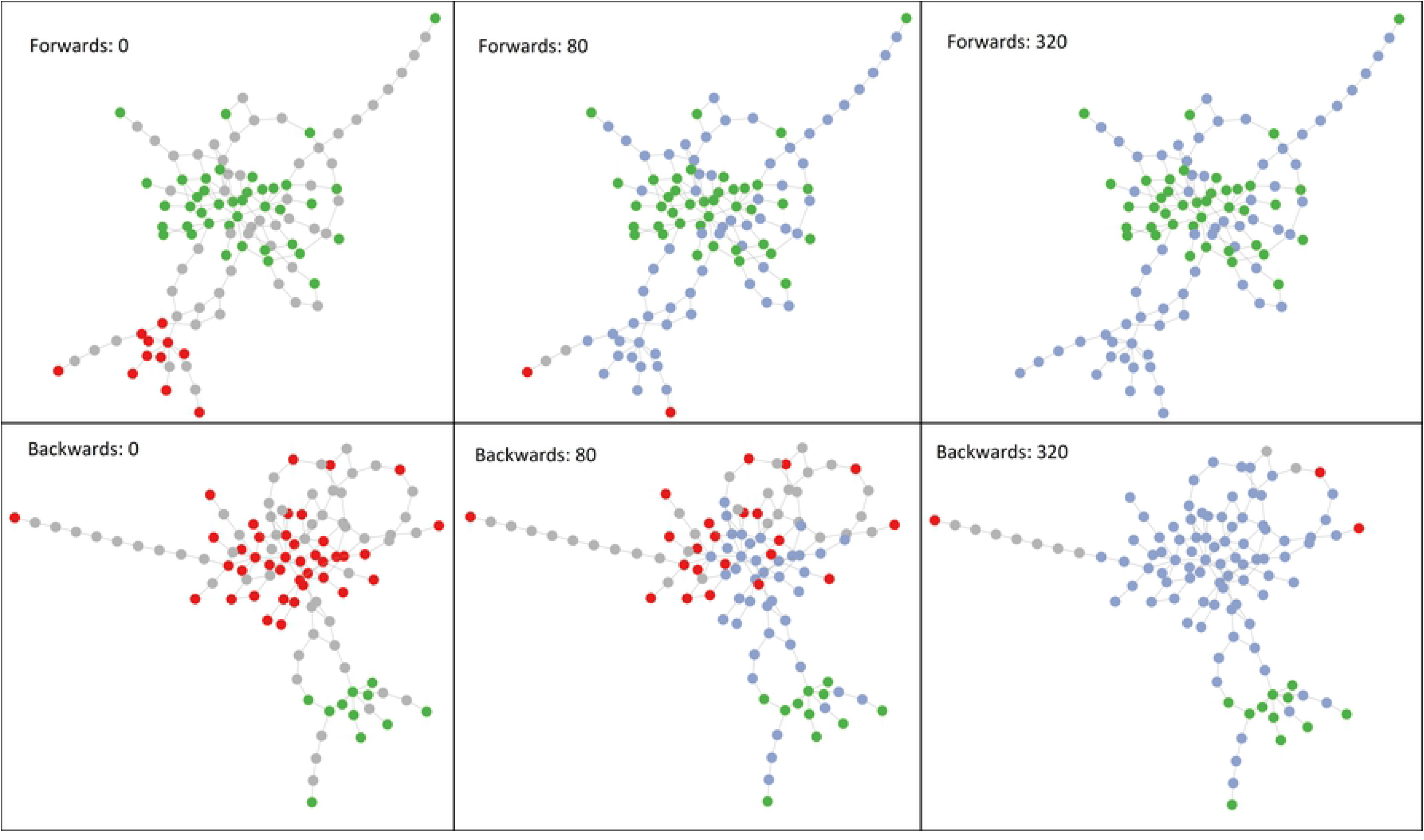
Visualization of evolutionary simulation. Dots represent unique variants. The simulation is a random walk that starts from the green potential infector variants with the gray intermediate variant dots and red potential infectee variant dots being unexplored. As the simulation carries out, dots turn blue as they are explored. The simulation is complete when all potential infectee variant dots are reached, or when all the red dots have turned blue. The top 3 frames show sample A evolving into sample B. The bottom 3 frames show sample B evolving into sample A. We can see that it is easier for A to evolve into B as it finishes faster, so we would estimate that A transmitted the virus to B rather than vice versa.

The model is described by the following system of equations:

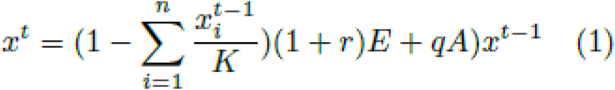

Here 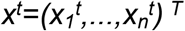 is a vector of abundances of haplotypes corresponding to the nodes of the graph *G* at time *t, K* is the maximum population size, *E* is an identity matrix and *A* is an adjacency matrix of *G*, and replication probability *r* and mutation probability *q* are defined as functions of the position-wise mutation rate ε and genome length *L*:

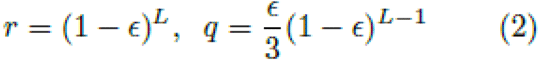

The evolutionary distance measure for a pair of populations *(i,j)* is defined as the time needed by the process from equation 1 to cover all nodes from the target *j* when starting only from the vertices of the potential infector *i*. This distance is considered to be infinite if the simulation takes longer than a threshold *T_max_*. The simulations are carried out for all pairs of patients and the calculated distances are stored in the distance matrix *M_T_*. Figure 3 depicts the relationship between this evolutionary simulation distance *T* and the minimum hamming distance between samples which has been shown to be a robust measure of relatedness [6].

**Figure 3:**
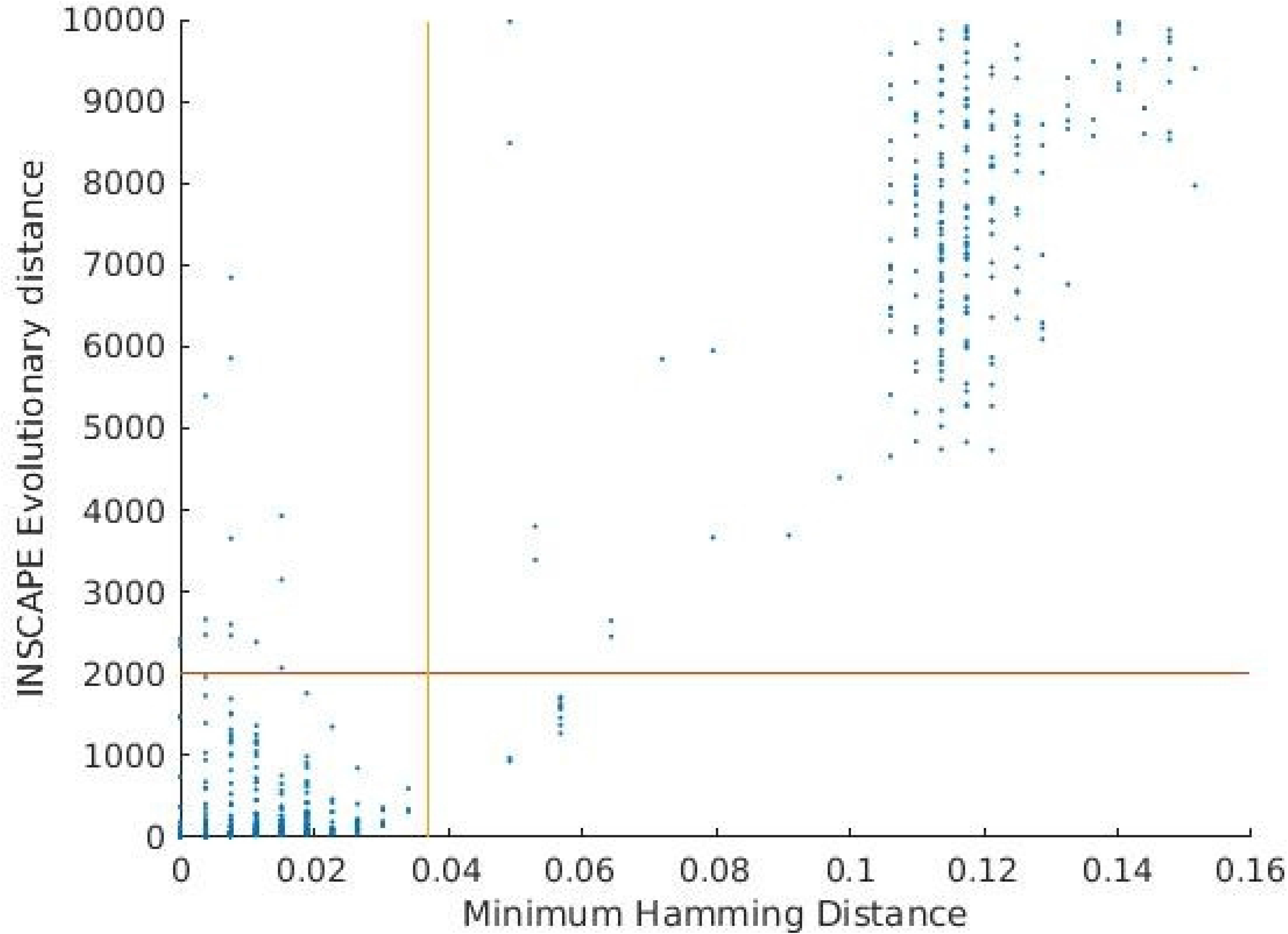
Distribution of PYCIVO distance measurements among our dataset. Most sample pairs fall either well below or above the empirically derived *T_max_* value of 2000. Evolutionary distances used in PYCIVO versus the minimum hamming distances between samples. The threshold for transmission is indicated by vertical and horizontal lines. Only 827 out of 5460 cases in our study were linked by the PYCIVO distance metric, 817 of these are also linked by minimum hamming distance.

The matrix *M_T_* is used to construct a directed *genetic relatedness graph, GT*, with nodes corresponding to patients, edges corresponding to pairs of patients with genetically related viral populations and their directions representing possible transmission directions. Using the minimal evolution principle, patients i and j are connected by an arc whenever *M_T_ [i,j] = min(M_T_ [i,j], M_T_ [j,i])* <∞.

If *G_T_* does not form a connected component with all members of the input, we cannot make any prediction about the primary case. A recent outbreak in which the entire transmission network was sampled should yield a connected component with the PYCIVO evolutionary distance metric [8]. Among all input in this component, the candidate primary case *a* is selected based on the parameter *V_1_(a)* defined as follows:

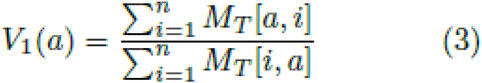

Where n is the number of samples present, the numerator represents the ‘in-evolution’ times, or the total time it took for all the samples to evolve to this one. The denominator represents the ‘out-evolution’ times, or the time it took for this sample to evolve into all the others. If this ratio is high, then that is indicative of a patient being a likely primary case candidate, as they were able to evolve to others more successfully than the other way around in simulations. If the ratio is low, that sample is not considered as a candidate for the primary case. For each sample, this value is the output for this portion of the algorithm. These values average to 1, with the best primary case candidates having higher values.

### Measures of Heterogeneity

The second method for primary case candidate determination within a transmission cluster works by selecting the infected individual with the highest amount of heterogeneity according to a custom index. We use several different measures to generate a composite score. Genetic heterogeneity has been shown to be higher for primary outbreak cases than other individuals sampled in an outbreak [6]. For each sample, 8 values are measured as shown below. *A* denotes the clinical sample of interest, where F(*A_i_*) and H(*A_i_,A_j_*) are shorthand for frequency and hamming distance, respectively. The chosen heterogeneity values are as follows:

1. Maximum intra-patient Hamming distance: the largest Hamming difference between two of the haplotypes

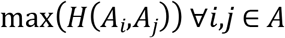
2. Mean intra-patient Hamming distance: the arithmetic mean of all the pairwise Hamming distance between pairs of sequences within the sample.

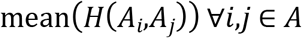
3. Nucleotide diversity: degree of polymorphism in a sample.

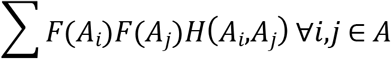
4. K-mer entropy: Shannon entropy over the set of 8-mers in the sample. Let A_k_={k_1_…,k_n_} be the set of k-mers.

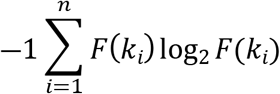
5. Haplotype entropy: Shannon entropy over the set of haplotypes in the sample.

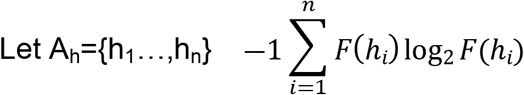
6. Average nucleotide entropy: Shannon entropy over the set of nucleotides in the sample. If we consider the alphabet N={A,T,C,G} and amplicon sequence length

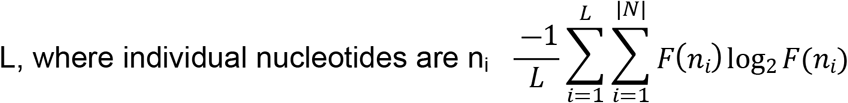
7. One step component entropy: Let A_0_={o_1_…o_n_} be the set of one-step hamming

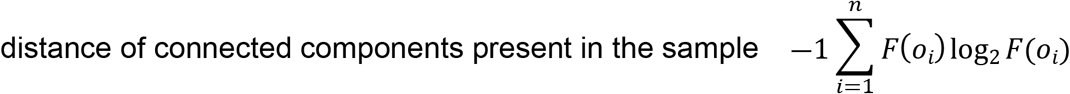
8. Mutation Frequency: Let M be the viral variant with the highest frequency

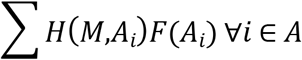

We chose these eight values because they were empirically shown to be higher for chronically infected individuals than acute ones, as shown in table 3. We combine them into a single scalar composite index for each sample. Using a matrix in which rows represent patients and columns represent each of the 8 features, each feature is ranked among patients in such a way that a more heterogeneous sample receives a lower rank. To obtain *V_2_* from this matrix, we compute the ranks in descending order along each index, then calculate the reciprocal of the harmonic mean of these ranks for each sample.

We choose this method rather than sorting in ascending order and using the arithmetic mean because we value a sample that is the most heterogeneous over most of the indices rather than being second or third among all indices. Using the reciprocal of the harmonic mean rewards samples which score the highest among a given index disproportionately.

### Implementation

PYCIVO uses data generated by the GHOST system, which processes genomic data generated by the Illumina MiSeq sequencer using several specially developed procedures [7]. There are some pre-processing requirements associated with this model. The input data should be a multiple sequence alignment in FASTA format, split into one file for each patient. Additionally, all reads in analysis must be same in length; length normalization can optionally be performed by PYCIVO using MAFFT with the -a option. Additionally, each file should not contain duplicate sequences. If there are two reads which have the same sequence, they are collapsed together into one haplotype with an associated frequency. This frequency should be at the end of the sequence ID following an underscore character.

## Results

PYCIVO was validated using data from 11 transmission clusters containing 105 samples and 1936 unique sequences [13-19]. In all cases, the primary case was epidemiologically identified. In addition, we used 10 modified transmission clusters in which the known primary case sample was removed in order to test PYCIVO specificity. One transmission cluster included only 2 cases and could not be used effectively as a modified cluster.

The cover times of simulated evolution from intra-host viral variants sampled from patient A to intra-host variants from patient B (out-evolution time) and from patient B to patient A (in-evolution time) are asymmetric; i.e., the time of evolution from A to B and from B to A are different. A good primary case candidate would have overall high in-evolution and low out-evolution times. This concept is illustrated in Fig. 2. In contrast, a simpler metric such as minimal or average hamming distance does not inform on which direction is more likely. The evolutionary time, however, correlates positively (*r=.65, p<10^-200^*) with hamming distances, as shown in Fig.3.

Both primary case indication values were calculated to issue primary case predictions for each outbreak. We aim to achieve accurate detection of the primary case only when one was actually sampled in an outbreak investigation. With this in mind, PYCIVO issues a “No Primary Case” (NPC) prediction when the primary case was not sampled or was removed from the data set.

This category of output is enabled by discrepancy between the two independent prediction methods, or when the genetic relatedness graph *G_T_* is not connected. In this case, PYCIVO informs the user that there is not a good candidate for the primary case within this group, claiming that they were likely not present in the samples given.

If neither of these occurred, then the z-score for the *V_1_* and *V_2_* vectors was used to determine the confidence of the prediction. We get two of these z-scores: one from *V_1_* and another from *V_2_*. Two levels of PYCIVO prediction were empirically established based off these z-scores: High Confidence (HC) when a sample has the highest value along both identification metrics, and both of their z-scores are above 2; Low Confidence (LC) when a sample has the highest value for both primary case indication values but one or both z-scores are below 2. Table 1 shows that LC predictions are much more common than NPC and HC predictions. This is an intentional feature of the software to give more veracity to NPC and HC predictions. This model is still in its early stages, so LC predictions at this point are prone to error and have little value without supporting epidemiological information.

**Table 1:**
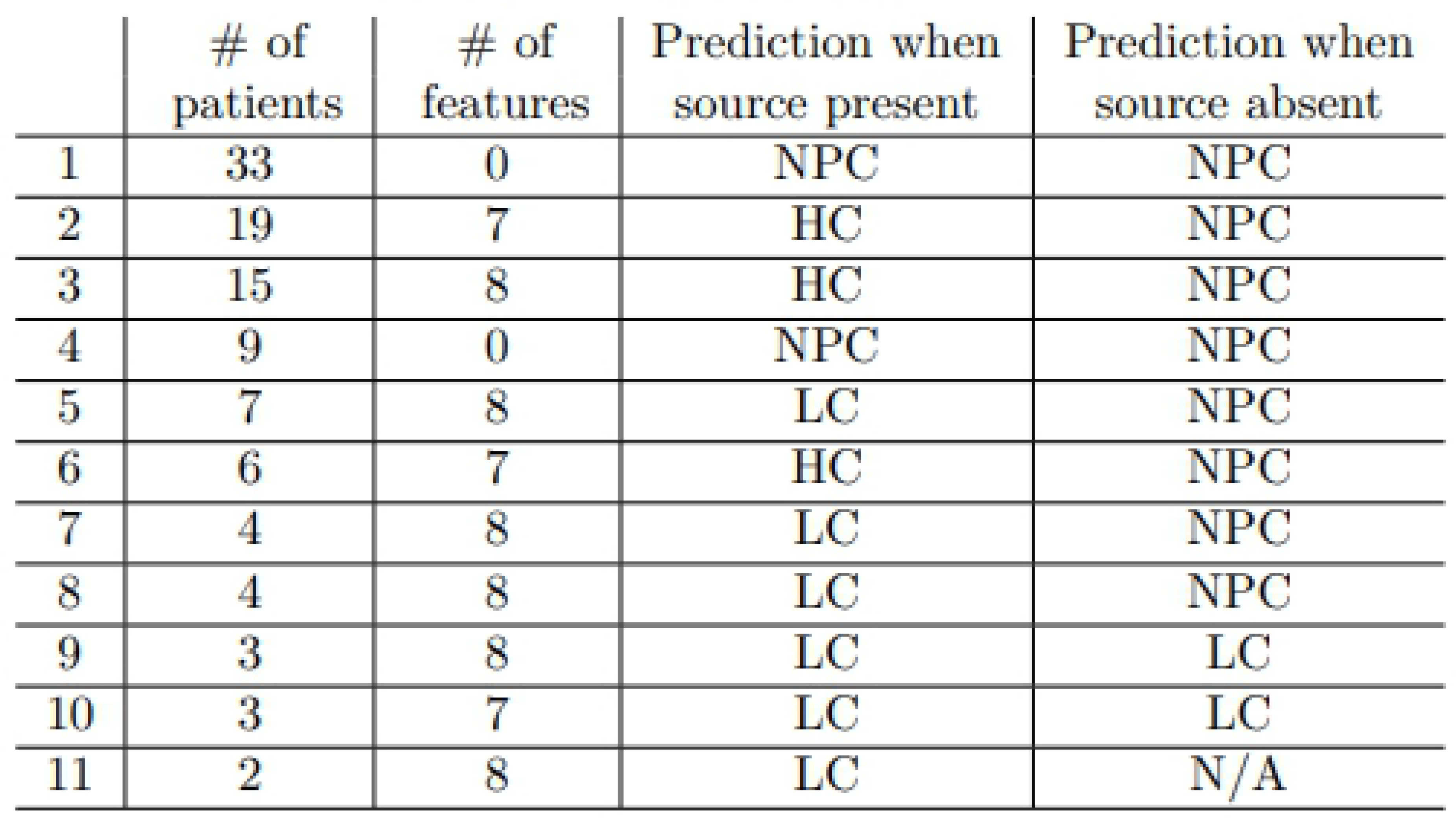
Outbreaks info

|  | # of patients | # of features | Prediction when source present | Prediction when source absent |
| --- | --- | --- | --- | --- |
| 1 | 33 | 0 | NPC | NPC |
| 2 | 19 | 7 | HC | NPC |
| 3 | 15 | 8 | HC | NPC |
| 4 | 9 | 0 | NPC | NPC |
| 5 | 7 | 8 | LC | NPC |
| 6 | 6 | 7 | HC | NPC |
| 7 | 4 | 8 | LC | NPC |
| 8 | 4 | 8 | LC | NPC |
| 9 | 3 | 8 | LC | LC |
| 10 | 3 | 7 | LC | LC |
| 11 | 2 | 8 | LC | N/A |

Tables 1 and 2 summarize the PYCIVO results obtained using the data from 11 known transmission clusters. The main results are as follows: (i) PYCIVO made primary case predictions in 9/11 clusters while the others were classified as NPC; (ii) HC predictions (n=3) were all accompanied by NPC when the primary case was removed; (iii) LC predictions (n=5) were accompanied by 3 NPC predictions as well as 2 erroneous LC predictions in the modified dataset, and the remaining outbreak did not have enough cases to conduct this analysis; (iv) the 2 cases in the modified dataset which issued LC predictions were outbreaks of only 2 patients; (v) NPC predictions were always accompanied by NPC predictions in the modified dataset.

**Table 2:**
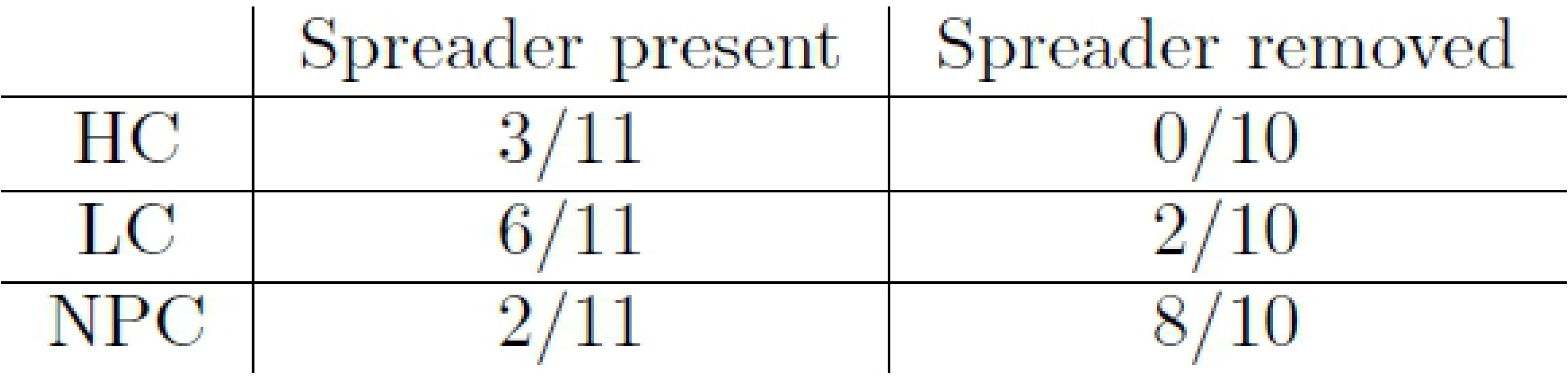
Outbreaks summary

Table 3 shows how each feature performed for the collection of outbreaks for which we had access. Many of the features show agreement with one another, except for the two indices at the Table bottom which were excluded due to their lack of empirical accuracy.

**Table 3:**
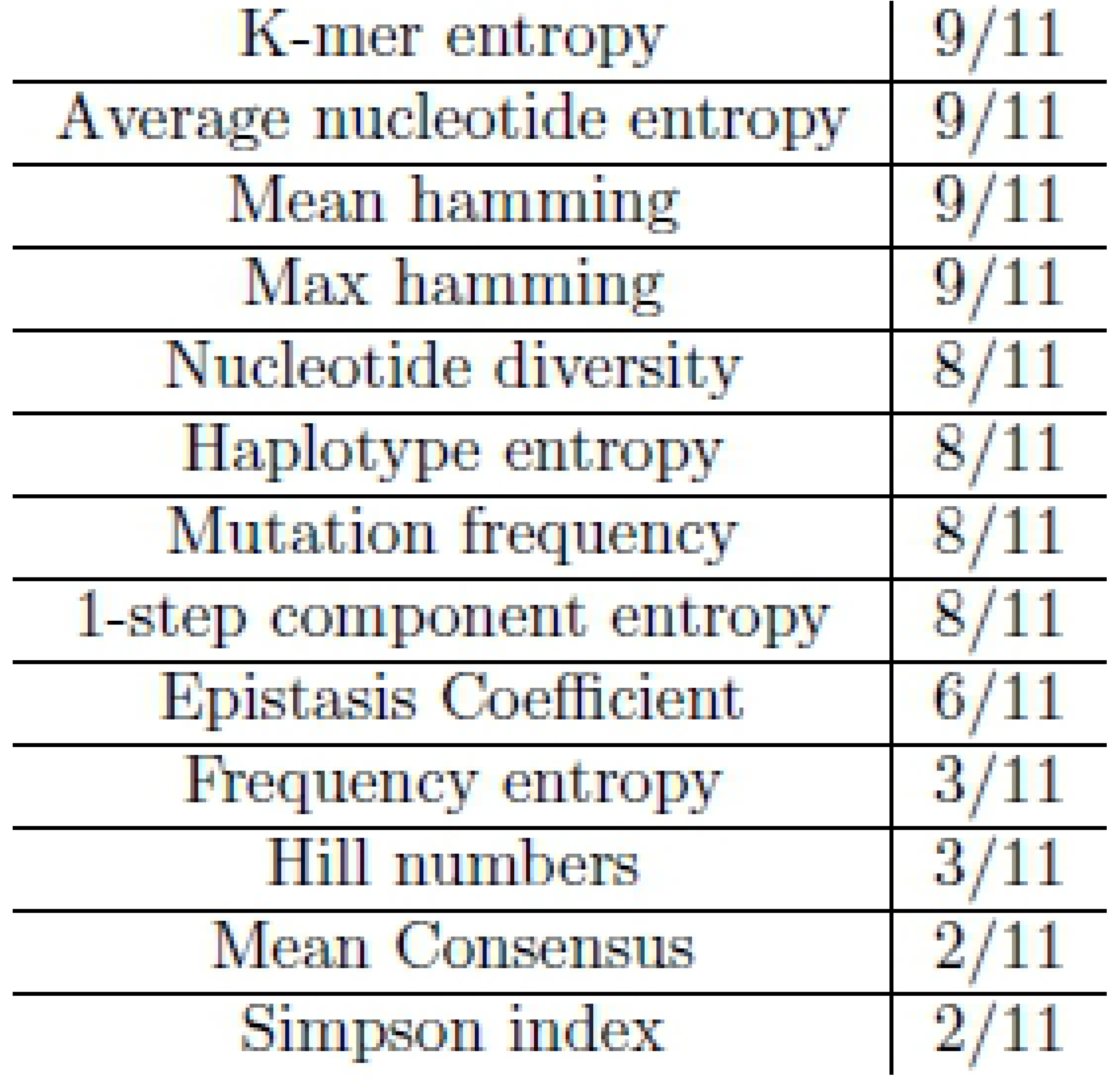
Feature Performance

|  |  |
| --- | --- |
| K-mer entropy | 9/11 |
| Average nucleotide entropy | 9/11 |
| Mean hamming | 9/11 |
| Max hamming | 9/11 |
| Nucleotide diversity | 8/11 |
| Haplotype entropy | 8/11 |
| Mutation frequency | 8/11 |
| 1-step component entropy | 8/11 |
| Epistasis Coefficient | 6/11 |
| Frequency entropy | 3/11 |
| Hill numbers | 3/11 |
| Mean Consensus | 2/11 |
| Simpson index | 2/11 |

Using this multitude of indices rather than just one of them produces a system which is more robust. This is so because artifacts in HCV evolution which reduce the primary case’s heterogeneity along one index will not be equally reflected in the other indices.

## Discussion

Genetic characterization of intra-host viral populations is often used for the detection of transmission clusters and for the investigation of outbreaks [6]. However, these genetic approaches, although highly effective in uncovering transmissions, are infrequently applied to tracking transmissions, owing to complexity of intra-host [20] and inter-host [5] viral evolution. Here, we present a new computational framework for the identification of the likely primary case in HCV outbreaks. The presented approach is based on simulation of evolution of intra-host HCV variants between cases involved in direct transmission during an outbreak, assessment of 2 identification values and statistical evaluation.

These methods enable PYCIVO to make the important NPC distinction, as well as preventing from making HC predictions on small outbreaks. This is so because the maximum z-score among a small set of random numbers will likely be less than 2, producing a LC prediction. It is difficult to determine the veracity of a source prediction for smaller outbreaks.

Three of the analyzed clusters are made up of three or less patients, which tend to produce LC predictions. In contrast, larger outbreaks make HC or NPC predictions more often. In agreement with this observation, there is a very high correlation between the minimum z-score used for prediction and the number of samples in the cluster (*r^2^=0.9322, p=2.4e^-5^*). This is advantageous for most outbreaks as majority of them have more than three patients, which steers PYCIVO towards the more conclusive NPC and HC predictions.

PYCIVO evaluates the strength of the molecular evidence available to identify the likely primary case of a recent transmission cluster. Our method considers the information provided by both inter-host and intra-host evolution, showing an accuracy of 81.8% when the primary case is present. The part of the method that deals with inter-host evolution uses the cover times of evolutionary simulations as an analog of evolutionary distance. If intra-host viral variants in one patient have higher out-evolution times, that means that they are unlikely to evolve to variants in other patients. If intra-host viral variants in a patient have higher in-evolution times, that means that they are unlikely to be evolved to. Intra-host variants of a likely primary case candidate should have a high in-evolution time and low out-evolution time. The part of the method that deals with intra-host evolution uses a composite heterogeneity score, which is generated via an ensemble of different metrics. The rationale for this second value is that, in general, HCV accumulates mutations during intra-host evolution and becomes more genetically heterogeneous over time [20]. Given that the primary case must have been infected for a longer time than all other incident cases, we can use this difference in duration of infection to infer the transmission direction. Indeed, our previous HCV analyses showed that the primary case is infected with a much more diverse HCV population than any incident case from the corresponding transmission cluster [5, 6]. This finding is supported with our earlier observation that, on average, the intra-host HVR1 nucleotide diversity is 1.8 times greater in patients with chronic than acute HCV infection [21].

Identification of the primary case is only possible with these methods if genetic samples from that patient are available. Otherwise, the incident case with the highest value may be erroneously classified as a source of infection in a transmission cluster. These problems of transmission-direction detection have been noted earlier [22, 23]. Therefore, our goal in developing this model was twofold: in addition to being able to predict the primary case, we aimed to be sensitive to input in which the primary case is neither known nor sampled. To achieve this goal, output was suppressed if there was uncertainty about who the primary case may have been, or if it appeared that they were not present among the samples. Output suppression thresholds were determined in via empirical thresholds outlined in the previous section.

PYCIVO is based on knowledge of intra-host and inter-host evolutionary dynamics of HCV shortly after a transmission event. Thus, the method’s performance will likely be less reliable if all the sampling times were not both close to the transmission event and close to each other. Over time, intra-host HCV populations from infected cases will both evolve away from one other and the population will grow more heterogeneous over time. The performance of this type of algorithm is impacted when one or more individuals experience superinfection, inflating the intra-host heterogeneity, which results in loss of association between the heterogeneity and duration of infection.

The previously published QUENTIN model [8] showed the same performance as PYCIVO when the transmission cluster does include the primary case. However, QUENTIN always chooses one sample among the input members as the primary case regardless of whether they are actually present in the input group of samples. In contrast, PYCIVO can issue the important NPC prediction on outbreaks, which do not have a clear primary case. The ability for PYCIVO to suppress erroneous output via this mechanism represents the major advantage of this model, as we cannot expect sequences from the primary case to always be available. Another difference is that in QUENTIN the network linking all the sequences of every pair of samples was a median-joining network rather than the k-step network used here, a change motivated by obtaining almost identical results with a considerably lower computational burden.

Our work has two major limitations. First, the datasets used in this study are limited in size. However, data on well-defined transmission clusters is exceedingly difficult to obtain. These data become available only after comprehensive epidemiological and molecular investigations. Second, the genomic data used here to devise and evaluate PYCIVO were not all obtained by deep sequencing but mainly by an older technology based on end-point limiting-dilution PCR and Sanger sequencing [20]. Both methods produce a population of variants, however, deep sequencing yields orders of magnitude more sequences. Although we previously found that statistical comparisons between these two methods on the same set of samples showed equivalent inter- and intra-host levels of heterogeneity [24], the use of PYCIVO with deep sequencing data needs to be further validated and updated. The results obtained in our study warrant research to further improve accuracy of the model to increase potential benefits of its application in the field. Utilization of differences in inter- and intra-host variability will likely continue as a core component of the model, but there are several avenues for modification in the future. The simplest way to improve upon the model is to accrue additional data from epidemiologically characterized outbreak investigations, which will either further validate the model or present opportunity to adjust parameters accordingly. We expect that our GHOST platform [7] developed to assist in identification of transmission clusters during outbreak investigation or similar technologies will provide ample data to improve the accuracy of this model in the future. Additionally, we plan further improvement of the model by updating the current strategy for calculating *V_2_* to include a feature vector based on predictions of infection duration by PHACELIA [25]. We may introduce new methodologies to the way in which *V_1_* is computed as well when novel data become available.

The epidemiological identification of outbreak sources is a very complex task. The genetic detection of a likely primary case can greatly facilitate investigation of outbreaks, assisting in identification of new cases and routes of transmission in specific epidemiological settings, and, thus, in guiding public health interventions for interruption and prevention of disease spread. However, it should be noted that although genetic testing can help detect the source of infection, it does not reveal the actual mechanism of transmission operating during outbreaks. Identification of such mechanisms and routes of transmission as well as the primary case role in an outbreak can be accomplished only through epidemiological investigation. For example, inadequate infection control, unsafe injection practices or drug diversion may be responsible for HCV transmission in healthcare settings rather than actions of source cases [26].

Understanding transmission of infections is crucial for effective public health interventions. The utility of transmission networks to public health interventions was demonstrated in simulation experiments [27-29]. Further improvement of accuracy will enhance potential benefits of the PYCIVO application to prevention and to development of more targeted interventions. However, genetic contact tracing and identification of primary cases present complex ethical issues associated with legal and social implications [30, 31]. Resolution of these issues is fundamentally related to data security. In this respect, it is important to note that the model reported here cannot collect, use, or produce personally identifiable information and is based exclusively on using genetic viral data.

## Conclusion

Here we present PYCIVO, a model that evaluates the strength of the molecular evidence available to identify the primary case for a transmission cluster. Our method takes into account the information provided by both inter-host and intra-host evolution, showing an accuracy of 81.8% when the primary case is present and with the important ability to issue ‘No Primary Case’ predictions in 805% of modified transmission clusters in which the epidemiologically verified primary case was removed.

## Disclaimer

The findings and conclusions in this report are those of the authors and do not necessarily represent the official opinion of the U.S. Centers for Disease Control and Prevention (CDC). Since PII is not being collected by CDC, output data from PYCIVO and GHOST cannot be used by CDC to identify or contact individuals.

## Acknowledgements

The authors would like to acknowledge Seth Sims for support in developing this software. The authors would additionally like to acknowledge Bailey Hannon, for proposing the name of the software. Funding: This work was partially supported by the National Institutes of Health grant 1R01EB025022-01.

